# Cryo-ET Reveals Distinct Gag Lattice Architectures in Virus-like Particles and Immature HIV-1

**DOI:** 10.64898/2025.12.08.693107

**Authors:** Benjamin Preece, Wiley Peppel, Haley Durden, Rodrigo Gallegos, Gabriel Clinger, Nicole Bohn, Antje Huwendiek-Poser, Gillian Ysassi, Allyson Roman, Giovanna Garcia, Tasheena Cly, David Belnap, Saveez Saffarian

## Abstract

HIV-1 is released from infected cells as immature virions whose membranes are supported by a Gag lattice. During maturation, this lattice is cleaved by the viral protease to release capsid proteins that assemble into the mature core. The architecture of the Gag lattice is central to this process, and the Gag lattice is targeted by maturation inhibitors that block cleavage. Using cryo-electron tomography, we compared Gag-only virus-like particles (VLPs) with immature HIV-1 virions and found that VLPs assemble denser and more complete lattices, exhibiting a strong correlation between lattice curvature and Gag copy number. In contrast, immature virions incorporate fewer Gag molecules and display weaker coupling between curvature and Gag stoichiometry. These findings show that while Gag alone can form the canonical immature lattice, additional viral components fine-tune lattice organization and curvature, potentially regulating protease accessibility, virion release, and the onset of HIV-1 maturation.

## Introduction

HIV/AIDS remains a major global health crisis, with approximately forty million individuals infected worldwide and seven hundred thousand people dying yearly from HIV-related causes worldwide^1^. Studying the Gag lattice in immature HIV-1 virions is crucial for understanding fundamental aspects of HIV maturation and developing effective antiviral therapies^2–4^.

As the primary structural component of newly assembled, immature virions, the Gag polyprotein forms a curved, hexagonal lattice at the plasma membrane of infected cells, driving virus assembly and budding^5–8^. Understanding the Gag lattice is vital because virion maturation, a prerequisite for infectivity, initiates from the lattice structure and involves the proteolytic cleavage of Gag and dramatic structural rearrangements within this lattice^9–11^. High-resolution structural studies, particularly using cryo-electron tomography (cryo-ET), have revealed key features like the six-helix bundle (6HB) at the CA-SP1 junction, which is suggested to act as a molecular switch regulating assembly and maturation^12–16^. The dynamic and incomplete nature of the Gag lattice, including its defects are believed to facilitate protease access and subsequent maturation^17–20^; furthermore, cellular cofactors like inositol hexakisphosphate (IP6) interact with and stabilize the immature Gag lattice and are fundamental for efficient assembly, maturation and infectivity^21–24^. Detailed knowledge of the immature Gag lattice provides a blueprint for designing new antiviral drugs that can interfere with HIV-1 assembly and maturation, such as maturation inhibitors that stabilize the lattice and prevent cleavage^2,25–28^.

The immature HIV-1 Gag lattice is notably incomplete, forming a contiguous but partial shell of Gag hexagons underlying the viral membrane. Cryo-electron tomography studies reveal that this lattice which consists of 250 to 500 Gag hexagons, covers only about 30-70% of the available space on the inner surface of the viral membrane^12,29^, in contrast to the estimated 5,000 Gag molecules required for a complete shell^30^. This incompleteness manifests as large gap-like defects and irregular imperfections, with these defects not attributed to disordered Gag molecules but rather to the absence of Gag in specific regions. Functionally, this incompleteness is critical: the resulting periphery of hexagons has Gag monomers with unfulfilled intermolecular contacts and provides more accessible targets for the viral protease^18,31^. The dynamic remodeling within the incomplete lattice is suggested to be essential for the dimerization and activation of the protease which is embedded in the Gag-Pol. Activation of HIV protease is a prerequisite for viral maturation and infectivity^17,32^.

The HIV-1 Gag polyprotein possesses the intrinsic ability to self-assemble into a lattice, forming virus-like particles (VLPs) in both cellular and in vitro environments, though the requirements and characteristics can vary between these settings.

In cells, expressing Gag protein by itself is generally sufficient to drive the formation and release of VLPs. These Gag VLPs often exhibit morphologies and sizes similar or even identical to immature HIV-1 virions when viewed by techniques like thin-section transmission electron microscopy (TEM)^33,34^. In vitro, purified monomeric Gag can assemble into VLPs, but often with specific additional requirements. Constructs of HIV-1 Gag lacking most of the matrix (MA) domain and the p6 domain (e.g., GagΔMA, ΔMA-CA-NC-SP2, or GagΔmyrΔp6) have been widely used to assemble membrane-less VLPs. This process is significantly influenced by the presence of nucleic acids, which are generally considered essential for efficient and correctly-sized membrane-less VLP formation^35–37^.

The Gag lattice in immature HIV virions differs from that in VLPs assembled in the same cells, primarily due to what is packaged within it. HIV virions specifically package the viral genomic RNA (gRNA) and the Gag-Pol polyprotein, while Gag VLPs package cellular RNA^38–40^. The RNA acts as a crucial scaffolding element for the immature Gag lattice, contributing to its stability and order through interactions with the Gag nucleocapsid (NC) domain^31,41–43^. The Gag-Pol polyprotein (at a low ratio to Gag^44^) is also incorporated into immature virions. This incorporation and subsequent precisely regulated activation of protease are critical for the proteolytic processing that drives the dramatic structural rearrangements leading to the mature, infectious virion^45,46^; In contrast, Gag VLPs, generally composed solely of Gag, lack the full genomic RNA and the functional Gag-Pol polyprotein, and therefore they do not undergo maturation.

During maturation, dramatic structural rearrangements occur, triggered by the proteolytic cleavage of the Gag and Gag-Pol polyproteins by the viral protease. This process transforms the virion from an “assembly mode” to an “infection mode” or “disassembly mode”^47,48^. Cryo-ET has been instrumental in capturing intermediate states within these transitions, revealing various maturation intermediates that represent distinct proteolytic and assembly stages^11^. The maturation pathway involves at least partial disassembly of the immature Gag lattice and the subsequent reassembly of the cleaved capsid protein (CA) into the characteristic mature conical core^11,47,49,50^ . Real-time monitoring techniques, such as NMR, allow observation of proteolytic cleavage events, like at the MA/CA junction, as they occur^51^. The intrinsic protein dynamics of CA and SP1 are critical for orderly maturation, and perturbations to these molecular motions by mutations or small molecules can inhibit the process^52^. Protease activation, which initiates these extensive changes, has been detected within producer cells prior to, or within seconds after, the release of free virions using sensitive techniques like instant structured illumination microscopy. Fluorescence lifetime imaging microscopy (FLIM) combined with single virus tracking has also revealed proteolytic cleavage of Gag *in situ* in a subset of particles, indicating that Gag processing can occur with a delay in many VLPs^53^. Beyond the core, the matrix (MA) protein also undergoes significant structural maturation, rearranging between distinct hexameric lattices and modifying the viral membrane by binding lipids, potentially by partially removing them from the lipid bilayer^54^. Moreover, super-resolution fluorescence microscopy (STED) has shown maturation-dependent redistribution and clustering of envelope (ENV) proteins on the viral surface, suggesting a dynamic coupling between internal structural rearrangements and changes on the viral exterior that prepare the virion for productive entry^49,55^. The transition from a rigid immature particle to a more flexible mature one is also a key dynamic change essential for efficient viral entry^49^. Furthermore, Gag dynamics within VLPs, including a measurable pseudo-diffusion rate, are sensitive to mutations within the SP1 region, suggesting its role in lattice ordering^32,56^. Experimental evidence highlights significant dynamics within HIV virions, both in their immature state and during the complex process of maturation.

To investigate the essential elements that hold the Gag lattice together and the inherent dynamics within the lattice, we utilized sub-tomogram averaging Cryo-ET to examine Gag VLPs and immature virions (NL4.3(D25N)) assembled and released from HEK293 cells. Our findings reveal that Gag VLPs incorporate more Gag and exhibit significantly enhanced lattice coverage (60-90%) when compared to NL4.3(D25N) virions which have a coverage of 30-60%. We further show that Gag lattice coverage in NL4.3( :(Δ 105-278 & Δ 301-332)(SS2-)(ΔENV))) VLPs, which retain HIV genomic RNA elements but lacks the packaging signal, proper ribosomal frameshifting to translate Gag-Pol, and also does not translate ENV, show similar increased Gag incorporation and enhanced lattice coverage (60-90%). Sub-tomogram averaging shows Gag CA-SP1 density at 6Å with similar structure resolved in Gag VLPs and NL4.3(D25N) virions. From these observations, we conclude that Gag-Gag interactions dominate the local structure of the Gag lattice while other viral factors, including Pol, Env and gRNA play a role in reducing dynamics as well as levels of Gag incorporation within HIV virions.

## Results

Comparing the Gag lattice in immature HIV-1 virions and Gag (VLPs) is particularly interesting because VLPs, formed primarily by the expression of the Gag polyprotein alone, are morphologically and structurally similar to authentic immature HIV-1 virions. This similarity makes VLPs a powerful model system for studying the fundamental mechanisms of virus assembly, as Gag itself plays the dominant role in defining the internal organization of the immature particle.

### Comparison of immature virions to Gag VLPs

Immature virions were harvested 48 hrs after transfection of pNL4.3:PR(D25N) in HEK293 cells. pNL4.3:PR(D25N) encodes a catalytically inactive viral protease due to the D25N substitution^57,58^ as shown in Figure S1. Virions were harvested and prepared for cryotomography according to protocols developed previously^59^ and described in methods. Cryo-specimens were imaged in a Titan Krios G3i transmission electron microscope (ThermoFisher) equipped with a Gatan BioQuantum K3 energy filter and direct electron detector. Tomograms were reconstructed from tiltseries acquisitions taken from −60 to 60 degrees as described in methods. Reconstructed tomograms show multiple immature virions in each tomogram as demonstrated in Figure 1A. These tomograms were then partitioned and analyzed using sub-tomogram averaging (STA) to visualize Gag hexamers as described in methods (Fig.1B). As shown in Figure 1, the Gag hexamers in NL4.3(D25N) virions cover only a fraction of the available space in the interior of the virion. In total we reconstructed 51, NL4.3(D25N) virions and found the Gag lattice in all virions to be contiguous with all Gag hexamers located within one large lattice continent within the virion as shown in Figure S2.

**Figure 1.**
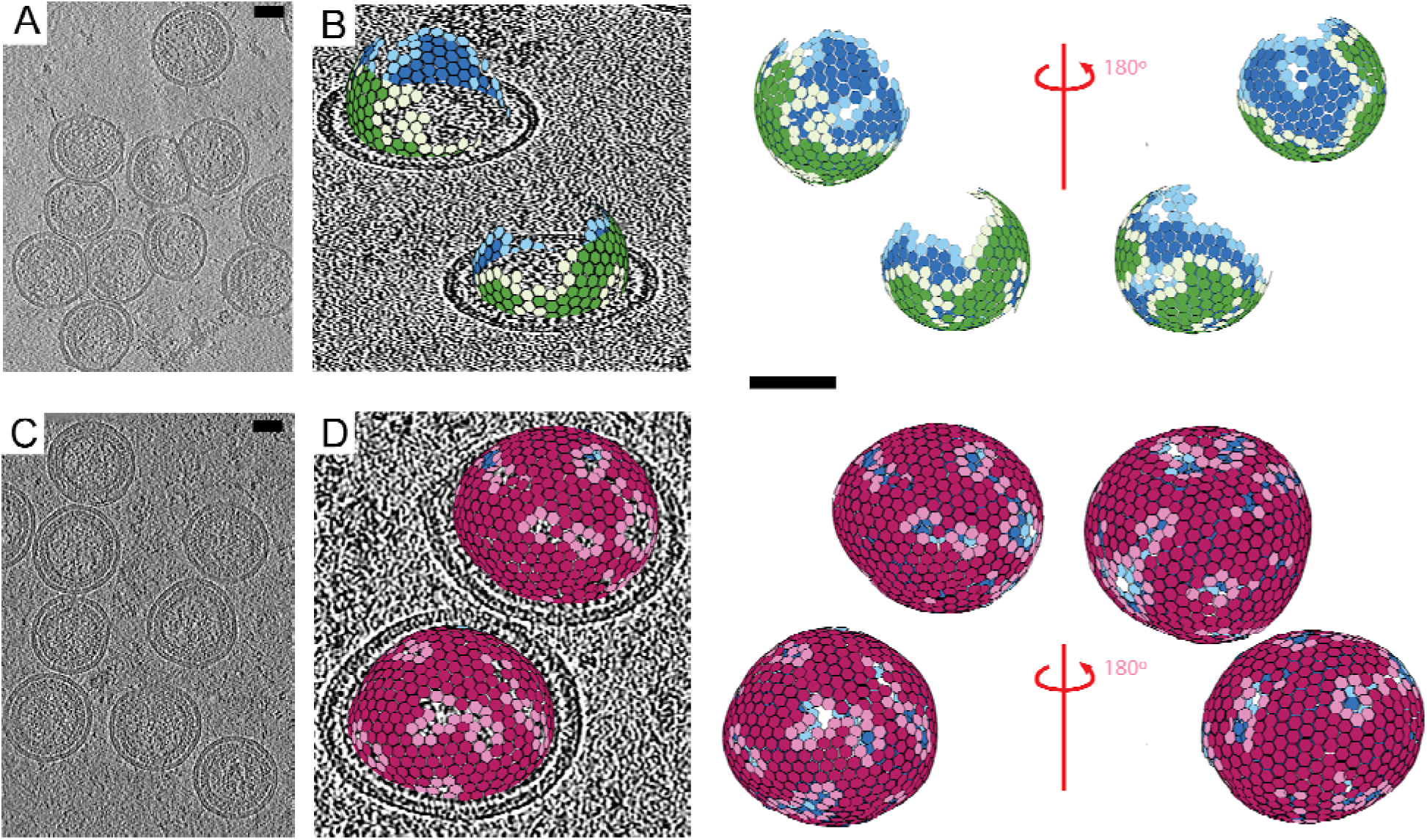
**(A)** Sample tomogram slice from NL4.3:PR(D25N). **(B)** Tomogram slices of NL4.3:PR(D25N) with hexamers fit to immature CA lattice positions. Immature Virions also shown without a tomographic slice and rotated 180° around the z axis. **(C)** Sample tomogram slice from humanized Gag VLPs. **(D)** Tomogram slice of humanized Gag VLPs with hexamer fit to immature CA lattice positions. VLPs are also shown without a tomographic slice and rotated 180° around the z axis. Scale bars are 50 nanometers in length.

We then utilized identical methodology to investigate Gag VLPs which were harvested 24 hours after transfecting a humanized Gag sequence, with identical amino acid sequence to the Gag from NL4.3 under a CMV promoter, into HEK293 cells as shown in Figure S1. Cryo specimens from Gag VLPs show particle rich tomograms of intact viral particles (Fig 1.C). Gag hexamer identified using STA, when compared with NL4.3(D25N) virions, show a significantly larger coverage of Gag hexamers within Gag VLPs. We reconstructed 44 Gag VLPs, and all VLPs had a single contiguous continent as shown in Figure S3.

We further verified the identification of lattice edges by averaging tomogram sections with one, two, or three neighbors missing along the edge of the lattice as shown in Figure 2A. To estimate coverage of the lattice, the lattice was projected onto a mesh surface fitted into the virion as shown in Figure 2B. These surfaces allowed estimation of both lattice coverage and radii of the virions/vlps as described in methods. Immature virions had an average of 2000 Gag monomers (SD +- 851 monomers) with an average CA lattice coverage of 57% (SD +-12%). These immature virions had an average radius of 70 nanometers (SD +- 5.5 nm) as shown in Figure 2C.

**Figure 2.**
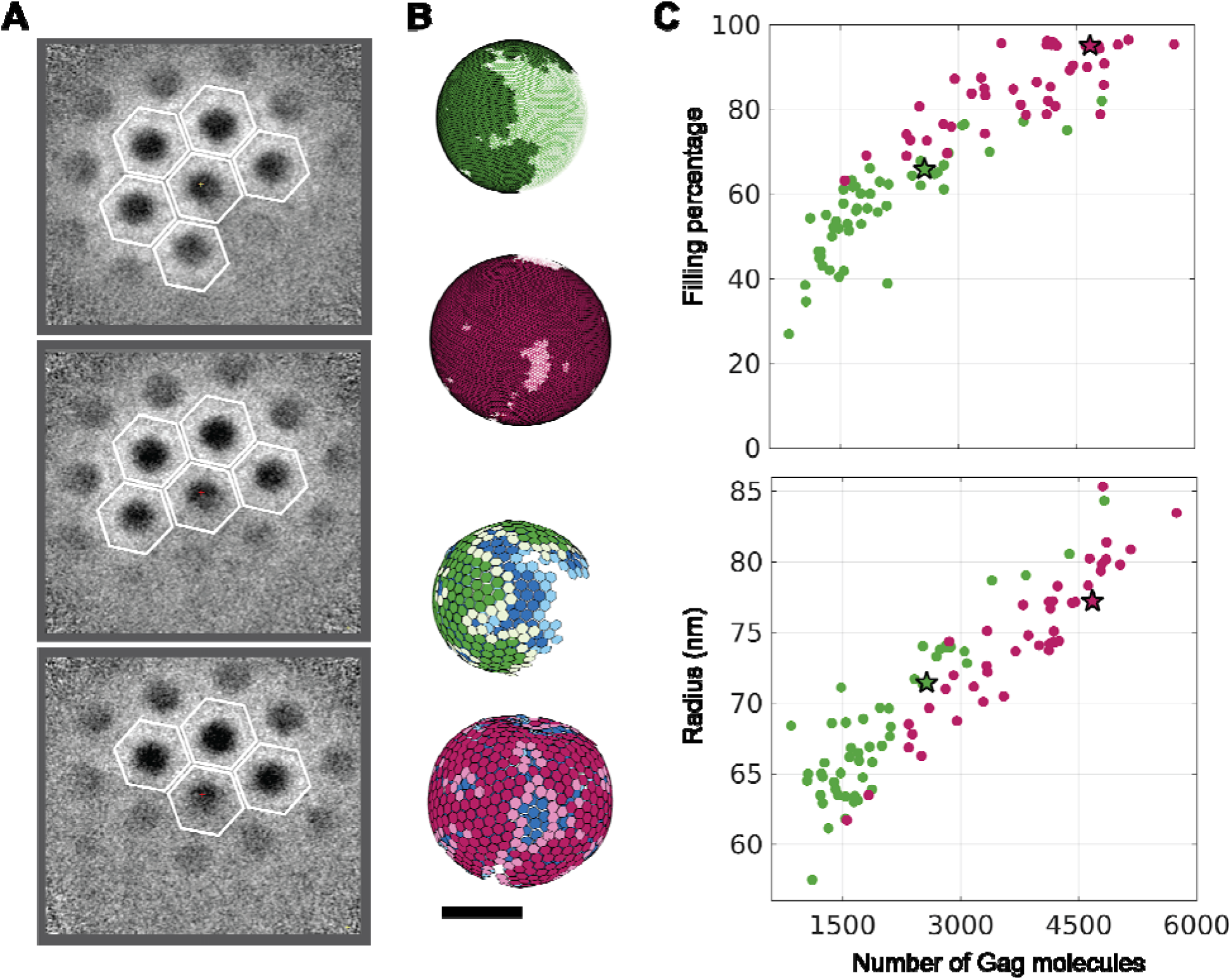
**(A)** Subtomogram averaged slices from immature lattice edge multireference STA. Frames highlight central Gag oligomer and adjacent hexamers with one, two, or three missing monomers in the central oligomer. **(B)** Immature CA lattice of NL4.3:PR(D25N) and hGag virions/VLPS shown with hexagons placed at subtomogram averaging CA hexamer centers of mass, and CA lattice coverage as shown fitted to an ellipsoidal surface. (hGag in Magenta and D25N in Green) **(C)** Graphs showing the number of Gag monomers per virion versus the immature lattice filling percentage and virion/VLP radius in nanometers. (D25N N=51 & hGag N=44) Scale Bar is 50 nm in length.

hGag VLPs had an average of 3800 Gag monomers (SD +- 960 monomers) with an average CA lattice coverage of 85% (SD +-9.5%). These hGag VLPs had an average radius of 75 nm (SD +-5.1 nm) as shown in Figure 2C.

A careful analysis of the quantification presented in figure 2 shows that the radius of both Gag VLPs and NL4.3(D25N) virions have a correlation with the number of Gag molecules within the particle; however, while the correlation of radius with the number of Gag molecules is very strong in Gag VLPs, the NL4.3(D25N) virions at equal number of Gag incorporation have a larger radii compared to Gag VLPs. As well, for NL4.3(D25N) virions incorporating less than 2000 Gag molecules, the radius and number of Gag molecules show far less correlation. In addition, Gag VLPs have a continuous distribution of Gag molecules within the VLPs while the number of Gag molecules within NL4.3(D25N) virions shows a bimodal distribution as shown in Figure S4.

Because the Gag translated in cells transfected with humanized plasmids under CMV promoters is unconstrained by the regulations of the genomic RNA during transcription, nuclear export and translation, we decided to also visualize Gag VLPs which were produced in cells in which Gag molecules were synthesized with regulations dictated by gRNA. To accomplish this, we mutagenized the NL4.3 backbone to remove the packaging signal and therefore reduce the probability of gRNA incorporation in the virions as previously described by Kutluay et al.^60^, to mutagenize the ribosomal slippage site as previously described by Garcia-Miranda et al.^61^ to abrogate translation of Gag-Pol, and to introduce a stop codon in ENV, which resulted in the NL4.3( :(Δ 105-278 & Δ 301-332)(SS2-)(ΔENV))) backbone shown in Figure S1 with mutations presented in Table S5.

Gag VLPs harvested 48 hours post transfection of the NL4.3( :(Δ 105-278 & Δ 301-332)(SS2-)(ΔENV))) backbone in HEK293 cells are shown in Figure 3A. These VLPs look identical to Gag VLPs generated by humanized Gag and have similar correlations between number of Gag molecules incorporated in the particles and their radius and lattice coverage as shown in Figure 3B.

**Figure 3.**
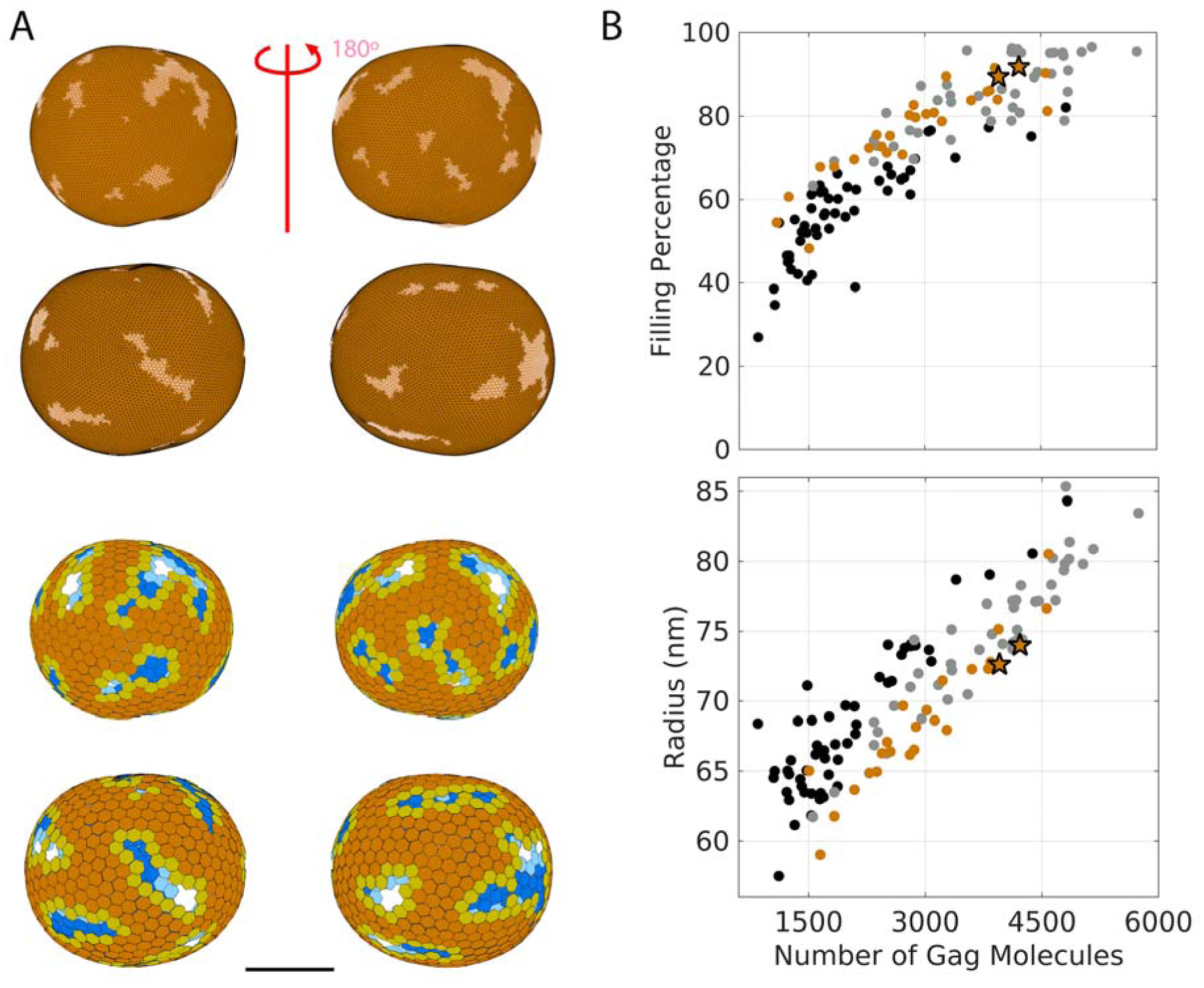
**(A)** Immature CA lattice mesh and hexagonal lattice map of NL4.3( :(Δ 105-278 & Δ 301-332)(SS2-)(ΔENV))) VLPs shown with a 180-degree z-axis rotation. All VLPs scale equally with a scale bar measuring 50 nm in length. **(B)** Graphs showing the number of Gag molecules per VLP versus their CA lattice filling percentage and VLP radius in nm. NL4.3( :(Δ 105-278 & Δ 301-332)(SS2-)(ΔENV))) shown in orange, NL4.3:PR(D25N) in black, and humanized Gag data in gray.

Robust subtomogram averaging was performed on both immature Virions and hGag VLPs to examine high resolution features of the CA lattices. Immature virion STA yielded a Gag-CA hexamer structure at 6.6 Å resolution, and hGag VLP STA yielded a 6.3 Å resolution structure. Both models were fitted using a rigid body model of PDB 5l93 for the CA-NTD region and PBD 5I4T for the CA-CTD-SP1 region, and no significant differences were noted as compared to the PDB models or between the density maps of the Gag average versus the immature virion average as presented in Figure 4.

**Figure 4.**
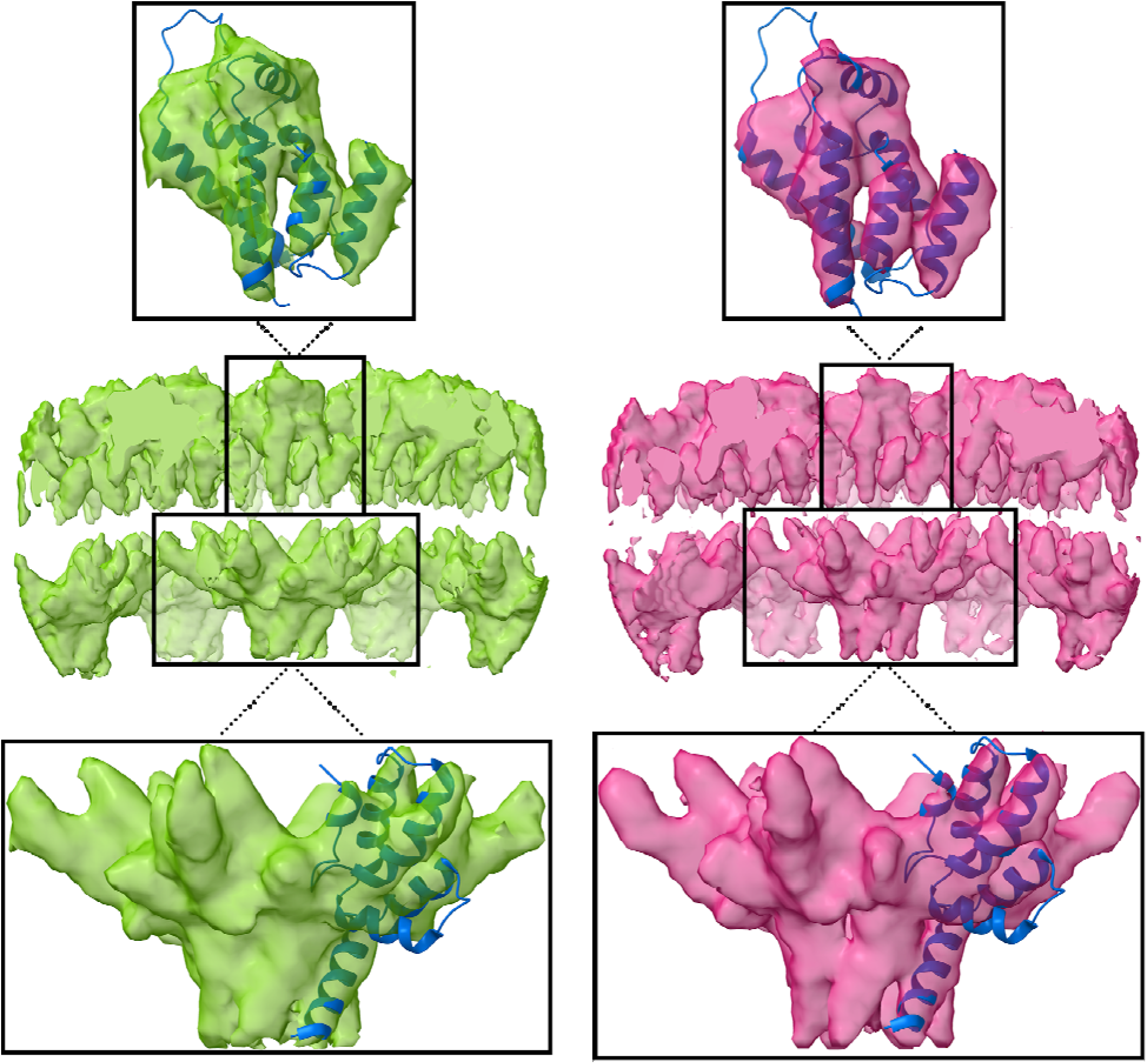
Subtomogram average of immature CA hexamers from NL4.3:PR(D25N), in green (6.6 A resolution), and hGag, in magenta (6.3 A resolution). Boxed regions show CA-NTD fitted to PDB 5l93 and CA-CTD-SP1 fitted to PBD 5I4T.

## Discussion

We showed that Gag virus-like particles (VLPs) have higher Gag incorporation and lattice coverage (60–90%) than immature HIV-1 virions (30–60%). Interestingly, Gag VLPs assembled by expressing humanized Gag constructs in HEK293 cells had identical structures to Gag VLPs assembled by expressing NL4.3( :(Δ 105-278 & Δ 301-332)(SS2-)(ΔENV))) in HEK293 cells. Below we will discuss these results in the context of HIV-1 assembly and release, mechanical properties of the Gag lattice and its impact in release and maturation, Gag-Pol:Gag-Pol association, and Gag-Pol auto-processing.

The more complete Gag lattice observed in the VLPs can provide a few insights regarding assembly and release of virions. It is plausible to suggest that the Gag lattice in the immature virions, recruits ESCRTs more efficiently and therefore results in release of virions before the virions are fully packed with Gag. One can imagine that such regulation can be useful because it would a) facilitate the release of virions prior to protease activation and b) create a significant budding scar with a lattice that has a substantial number for Gag molecules at the edge of the lattice, which is suggested to be important for protease access. Regardless of the advantages, such a mechanism would suggest that other factors aside from Gag are essential for ESCRT recruitment. This suggestion is supported by the following observations: It was suggested that the action of the cellular Endosomal Sorting Complexes Required for Transport (ESCRT) machinery, which mediate membrane scission and release of the immature virions, leaves a significant gap or “budding scar” in the spherical Gag shell^29^, accounting for the 30-60% of coverage of the Gag lattice in these virions. ESCRTs are recruited through their interactions with either the p6 domain of Gag, Gag NC, or indirectly through interactions with ubiquitin associated with Gag, all of which are dependent on the Gag protein^7,62^. If the budding scar was a result of ESCRT interactions, it would be reasonable to predict that such a budding scar should exist in both immature virions and Gag VLPs. It is therefore plausible to suggest that not every interaction of ESCRTs with the releasing virions would result in a budding scar. This observation would also agree with the Gag lattice observed in the released HIV-2 Gag VLPs, which did not report to have a major “budding scar”^63^. Experiments monitoring the release of HIV virions with abrogated ESCRT interactions showed that interaction of ESCRTs with HIV-1 are more nuanced than just facilitating the release of virions. The timing of HIV release also seems to be important for release of infectious virions, as a delay in release resulted in protease activation before release, which caused loss of essential viral enzymes and release of non-infectious virions^64^. While our data shows that the radius of the virion/VLP generally correlates with the number of Gag molecules incorporated in the virion/VLP, there is a distinction between this correlation in Gag VLPs where the correlation appears more tight and exact compared to immature virions; immature virions appear to be clustered into two sizes, with virions having 1000-2000 Gag molecules and a diameter of ∼65 nm and virions between 2000-3000 molecules having a radius of ∼75 nm. From these observations, one can project that there seems to be a preset curvature which corresponds to a virion with a 65 nm radius. As more Gag molecules are recruited to the virion, the lattice is filled along this predetermined curvature, and the virion is released before there is a chance to grow the lattice above 60% of the inner surface of the virion. In contrast, during Gag VLP assembly, the VLP grows in diameter as more Gag molecules incorporate within the VLP and eventually release with a large fraction of the inner surface of the virion filled with the Gag lattice and the radius of the VLP directly proportional to its Gag content. This correlation would argue that there is no strict Gag curvature and the curvature of the VLP is adjusted based on the number of Gag molecules incorporated. It remains to be seen by follow up experiments if this suggestion is valid, and if so, which of the HIV cofactors not present in the VLPs is responsible for maintaining the curvature of the immature virions.

One element absent in Gag VLPs is the HIV gRNA. gRNA is not merely encapsulated but is an active participant in initiating Gag multimerization, scaffolding the immature virion, influencing the order and stability of the Gag lattice, and coordinating protein-RNA interactions throughout the assembly process. It has been shown that gRNA plays a structural role in the immature HIV virions^65^ with two copies of gRNA being selectively incorporated into each virion during assembly^66–68^. Live imaging experiments show the recruitment of gRNA as an early event in immature virion assembly^69^ with Gag-gRNA interactions observed within the cytosol prior to initiation of assembly^70^. Specific recognition of the viral RNA packaging signal ( ) by Gag promotes the nucleation of Gag-Gag interactions at the early stages of immature viral particle assembly^71^ with Gag multimerization on viral RNA as a critical property required for efficient genome packaging^40^. Beyond nucleation, it is shown that gRNA and the membrane are critical constituents that actively promote Gag multimerization through scaffolding^31^. Gag’s extension and conformational changes are contingent on simultaneous interactions with both the membrane and nucleic acids. These interactions within the CA-SP1 region trigger conformational changes to the six helix bundle, acting as a switch for assembly^13,72^. Specifically, crosslinking-immunoprecipitation (CLIP) sequencing of the viral gRNA in immature virions shows extensive contact between Gag and gRNA through the full length of the gRNA in the immature HIV virions^60^. It is therefore plausible to propose that the extensive interactions of gRNA with the Gag lattice in immature virions would result in additional structural rigidity of the Gag lattice in immature virions when compared to Gag VLPs, which package only cellular RNA. Why cellular RNA would provide less stability compared to gRNA is unclear.

Having a more rigid Gag lattice in immature HIV virions can also be critical for Gag-Pol auto-processing. Some models of Gag-Pol auto-processing suggest that Gag-Pol:Gag-Pol association is critical for the initial dimerization of the protease and initiation of the Gag-Pol auto-processing. It is reasonable to assume that a more rigid Gag lattice will trap the Gag-Pol molecules and inhibit their rapid movement and association with one another. This would be an important mechanism which would safeguard HIV enzymes from premature proteolysis and subsequent protein loss to the cytosol of the infected cells. We currently do not know where Gag-Pol proteins are within the Gag lattice of the immature virions, therefore creating a detailed model of Gag-Pol auto-processing is beyond our reach.

Taken together, our data provide strong evidence for an unknown factor modulating Gag incorporation into the Gag lattice in immature HIV-1 virions. Further investigation is warranted to investigate Gag incorporation and the role of the dynamic movement of the immature Gag lattice in initiating viral maturation.

## Materials and methods

### Cell Plating and Transfections

Human embryonic kidney HEK293 cells were cultured in T-25 flasks using TrypLE Express Enzyme (Gibco, Thermo Fisher Scientific, Waltham, MA, USA) and Dulbecco’s Modified Eagle Medium (DMEM) supplemented with 4 mM L-Glutamine, 4.5 g/L Glucose, sodium pyruvate (Cytiva, Marlborough, MA, USA), and 10% fetal bovine serum (Gibco). Cells were incubated at 37 °C in a humidified atmosphere containing 95% air and 5% CO2 and passaged every other day or upon reaching confluency. Once the HEK293 cells reached 70–90% confluency, they were seeded onto 10 cm culture dishes at 9 mL of medium per dish and incubated for 24 h prior to transfection. At approximately 60% confluency, cells were transfected with the designated plasmid. Unless otherwise noted, transient transfections were performed using Lipofectamine 2000 reagent (Life Technology, Carlsbad, CA, USA). For each transfection, 20 µg of plasmid DNA and 40 µL of Lipofectamine 2000 were separately diluted in 300 µL of Opti-MEM (Gibco) and incubated at room temperature for 5 min. The two solutions were then combined and allowed to incubate for an additional 20 min at room temperature to form DNA-Lipofectamine 2000 complexes. These complexes were added dropwise to each culture dish containing cells at 60% confluency. Cells were then incubated at 37 °C for 48 h before harvesting.

### pNL4.3 Plasmid Preps and Mutagenesis

Cloning and plasmids: Proviral clone pNL4.3 was obtained through the NIH HIV Reagent Program, Division of AIDS, NIAID, NIH: (HIV-1), Strain NL4-3 Infectious Molecular Clone (pNL4-3), ARP-114, contributed by Dr. M. Martin.

DNA amplification and purification for transfection: A measure of 4 μg of lyophilized plasmid was resuspended in 10 μL of DNase/RNase-free distilled water and stored at −20 °C. Following this, 1 μL (0.4 μg) of each plasmid was added to 10 μL of Invitrogen MAXEfficiency chemically competent DH5α cells. Plasmid and cells were incubated on ice for 30 min and then heat shocked at 42 °C for 40 s. This was followed by a 2 min recovery on ice. Then, 250 μL of room-temperature LB was added to each tube and then plated on LB-agar plates with 0.01% ampicillin. Plates were inverted and incubated overnight at 37 °C. A single colony was used to inoculate 5 mL of LB with 10 ug/mL of ampicillin and incubated for 8 h at 37 °C whilst being shaken at 225 rpm. Then, 1 mL of this starter culture was inoculated into 500 mL liquid LB with 20 μg/mL of ampicillin and cultured overnight in a shaker (225 rpm) at 37 °C. DNA for transfections was purified per manufacturer’s protocol, using the GeneJET Plasmid Maxiprep Kit (Thermo Fisher Scientific, Waltham, MA, USA), and eluted in 1 mL of elution buffer.

### Preparation of OptiPrep™—Iodixanol Step Gradients

Two solutions (A and B) are prepared to make the step gradient. Solution A is made from 30 mL of 100 mM Hepes and 70 mL Phosphate-Buffered Saline (PBS). Solution B consists of 10 mL of 100 mM Hepes and 90 mL PBS. Both of these solutions are then filtered through 0.22 μm filters (CellTreat Product Code: 228747) for sterilization. Then, 10 mL of Solution A is mixed with 20 mL of OptiPrep™ Density Gradient Medium from Sigma Aldrich. This creates a solution of 40% OptiPrep™ and will be used as the bottom step in the gradient. For the top step, mix 7.5 mL of the 40% OptiPrep™ solution into 12.5 mL of Solution B to make a 15% OptiPrep™ solution.

### Preparation of 10X 10 nm Gold Beads

For use with cryo-electron tomography, functionalized gold nanoparticles must be added to the resuspended pellet at the end of the purification process. 1 mL of 10 nm BSA Gold Tracers are concentrated by spinning in a 100 kD Millipore Amicon Ultra, for five minutes at 5000 rpm. Then, 1 mL of STE Buffer (20 mM TRIS-HCl, 100 mM NaCl, 1 mM EDTA, pH: 7.4) is added and spun again for 3 min. The desired final concentration is 10-fold over stock.

### Freezing of Virions on EM Grids

Immediately following each VLP harvest, a one-to-one volume fraction of concentrated 10 nm gold fiducials were added to the resuspension. Then, 3.5 uL of the resulting sample was added to glow-discharged ultrathin (2 nm) carbon on Quantifoil R2/1 holey carbon film on 200 Cu mesh EM Grids (Ted Pella, Redding, CA, USA; Quantifoil Micro Tools GmbH, Großlöbichau, Germany). Following a single 2 second blot and a 1 minute wait, samples were plunge-frozen in a liquid ethane/propane mix, using a Vitrobot Mark IV (Thermo Fisher Scientific), with the incubation chamber set at 4 °C and 90–100% humidity.

### Cryo-Electron Tomography of Viral Particles Embedded in Vitreous Ice

Next, cryo-specimens were imaged in a Titan Krios G3i transmission electron microscope (ThermoFisher) equipped with a Gatan BioQuantum K3 energy filter and direct electron detector. Tilt series from −60° to +60° were recorded at 3° steps over holes in the carbon film via SerialEM software, version (p. 5) (University of Colorado, Boulder, CO 80309, USA) [55]. The microscope was operated at 300 kV, with images having a pixel size corresponding to 1.4 Å^2 at the specimen. The slit width of the energy filter was 20–30 eV. Tilt series were recorded at a target defocus between −5 µm to −1 µm. The total electron dose at the specimen was 100–125 electrons per square Å.

### Tomogram Reconstruction

To generate tilt series stacks from the raw micrographs, 4–10 frames per tilt angle were automatically aligned using SerialEM. The resulting 41-frame tilt series were aligned using 10–20 gold fiducials per tilt series in IMOD (University of Colorado, Boulder, CO 80309, USA). Using IMOD, each tilt series was then fitted for CTF correction, using Ctfplotter, and dose filtered at 80% of standard. Tomograms were then 3D CTF corrected and reconstructed in 15 nm slabs for final reconstructions of 1000–1800 slices or ∼140–250 nm total thickness.

### Sub-tomogram Analysis

Subtomograms were extracted from full tomograms by estimating VLP/virion centers and CA lattice radii by hand and oversampling positions along a resultant sphere^73^. Subtomograms were iteratively aligned using Dynamo STA software^74^. Subtomogram averages were subboxed, with a new subtomogram located at each identified CA hexameric center. Subtomograms were eliminated based on standard CA center to center euclidean distance, and cross-correlation score. These new centers were re-cropped as new subtomograms from the original full tomograms and subject to additional rounds of refined alignment in Dynamo. The aligned positions were extracted from Dynamo, and likely CA hexamers with fewer than 3 neighboring hexamers were also eliminated. Resulting CA hexamer positions were visualized in 3D, and centers not following the contiguous lattice were eliminated. These final CA hexamer positions and their euler rotations were used to template the lattice with hexagons as shown in the figures. For high resolution subtomogram averages, the final Dynamo alignment tables were subject to strong Cross-Correlation thresholding and split for Gold Standard alignment and resolution estimation. Following further refined alignment in Dynamo for the split data sets, Relion 4 was utilized to estimate FSC curves for resolution estimation.

### Mesh Fitting for Radii and Coverage Calculations

CA lattice center of mass coordinates for each virion/VLP were fit to an ellipsoid (Yury (2025). Ellipsoid fit , MATLAB Central File Exchange). Using a custom Matlab script, a mesh of ∼10000 vertices was then constructed from the ellipsoid fitting results and subjected to iterative laplacian fitting and smoothing to bring lattice curvature in line with the CA lattice coordinates. Mesh positions were then identified that would be covered by protein density and marked as such. Total vertices versus protein density covered vertices were compared to extract immature Gag lattice coverage percentages. Meshes were fit again to ellipsoids and ellipsoidal semi-axis averaged to estimate the CA-lattice radii. Average CA lattice distance to viral membranes was calculated from STA structures and added to CA-lattice radii to estimate virion/VLP radii.

### Edge Detection and Multi-Reference Alignment

Gag oligomers with fewer than 6 neighbors, based on euclidean center-to-center measurements, were identified using a custom matlab script as edges to the immature Gag lattice. These edge oligomers were subjected to Multi-Reference alignment (MRA) in Dynamo using synthetically derived templates. Templates of oligomers missing one, two, or three neighboring Gag hexamers were created by masking the high resolution Gag structure from standard STA to reduce protein density in one, two, or three of the neighboring Gag domains.

### Author Contributions

Conceptualization, B.P., W.P., and S.S.; methodology, B.P., W.P., H.D., D.B. and S.S.; software, B.P., and S.S.; validation, B.P., W.P. and S.S.; formal analysis, B.P. and S.S.; investigation, B.P., W.P., H.D., R.G., G.C., N.B., A.H., T.C., G.Y., A.R., G.G., D.B., and S.S.; resources, S.S.; data curation, B.P. and S.S.; writing—original draft preparation, B.P., W.P., H.D., R.G., G.C., N.B., A.H., T.C., G.Y., A.R., G.G., D.B., and S.S.; writing—review and editing, B.P., W.P., H.D., R.G., G.C., N.B., A.H., T.C., G.Y., A.R., G.G., D.B., and S.S.; visualization, B.P. and S.S.; supervision, B.P. and S.S.; project administration, B.P., H.D., G.Y. and S.S.; funding acquisition, S.S. All authors have read and agreed to the published version of the manuscript.

## Acknowledgments

This research was funded by National Institutes of Health (NIH) R56AI150474-06A1 and NIH R01AI186663. The following reagent was obtained through the NIH HIV Reagent Program, Division of AIDS, NIAID, NIH: TZM-bl Cells, ARP-8129, contributed by Dr. John C. Kappes, Dr. Xiaoyun Wu, and Tranzyme Inc.

