## Supplemental figures and text for "Cryo-ET Reveals Distinct Gag Lattice Architectures in Virus-like Particles and Immature HIV-1"

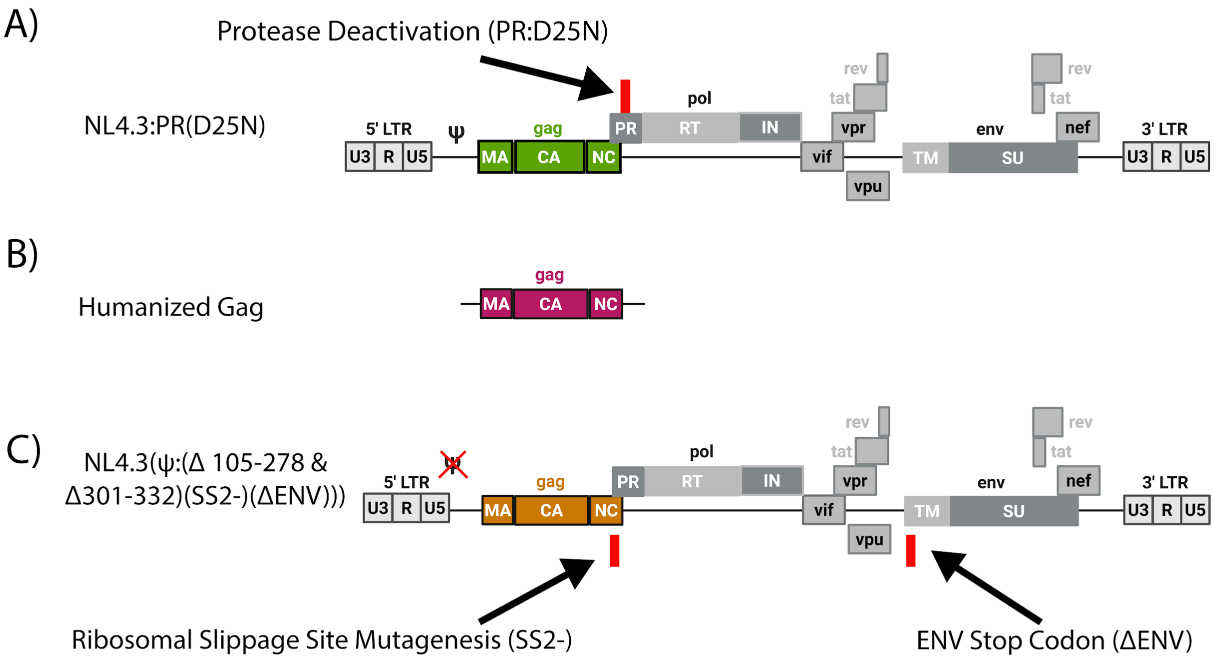


**Figure S1. (A)** Schematic representation of NL4.3:PR(D25N) genome. **(B)** Schematic representation of humanized Gag genome. **(C)** Schematic representation of NL4.3(Ψ:(Δ 105-278 &Δ301-332)(SS2-:((c.2085T>C,c.2088T>C,c.2130T>C, c.2133T>C))(ΔENV:(c.6403_6405del)))).

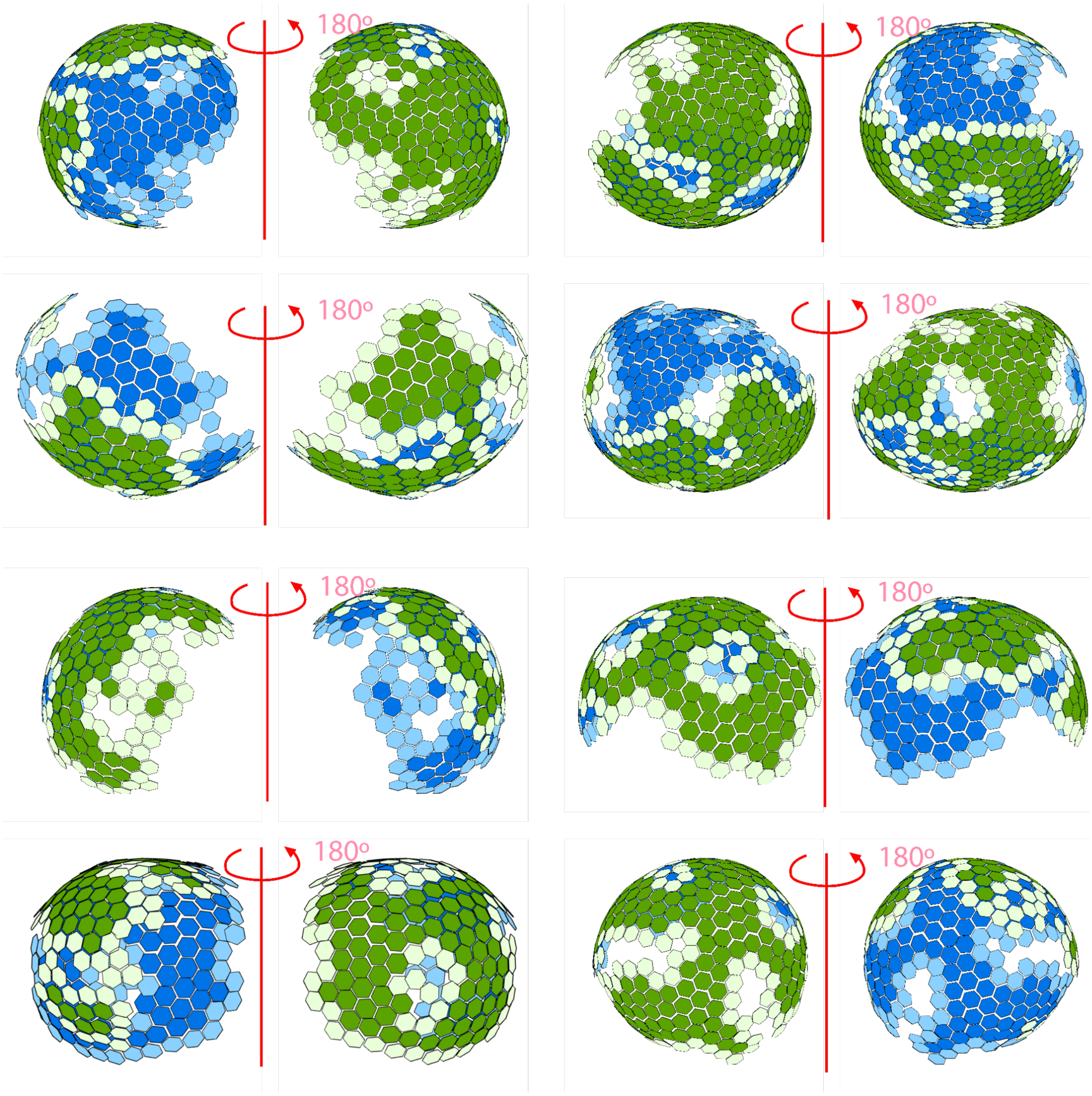


**Figure S2.** Gallery of NL4.3:PR(D25N) immature virions shown with hexagons placed at CA lattice centers of mass as calculated via STA. Immature Virions are also shown rotated 180 degrees around the z axis.

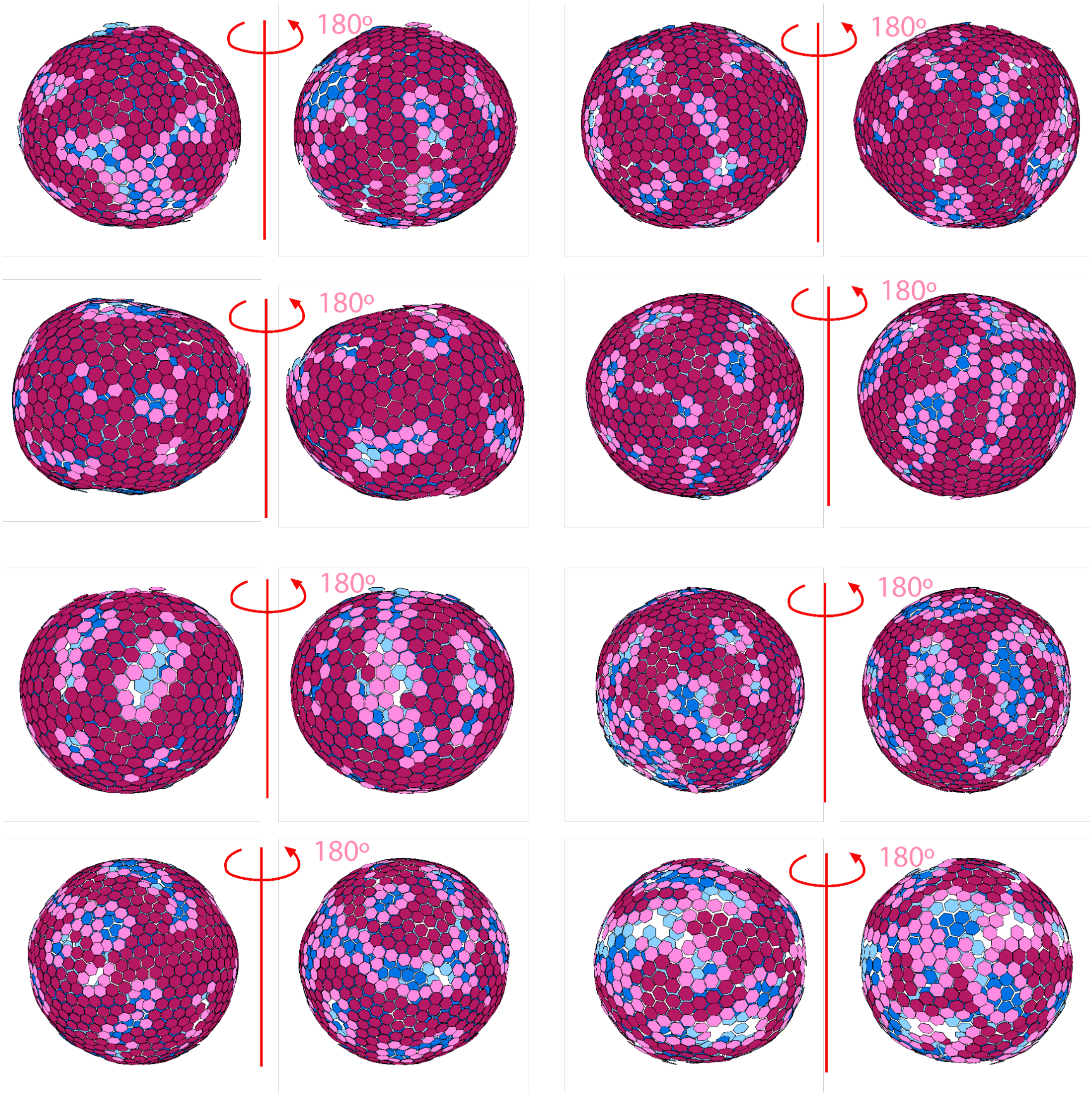


**Figure S3.** Gallery of humanized Gag VLPs shown with hexagons placed at CA lattice centers of mass as calculated via STA. VLPs are also shown rotated 180 degrees around the z axis.

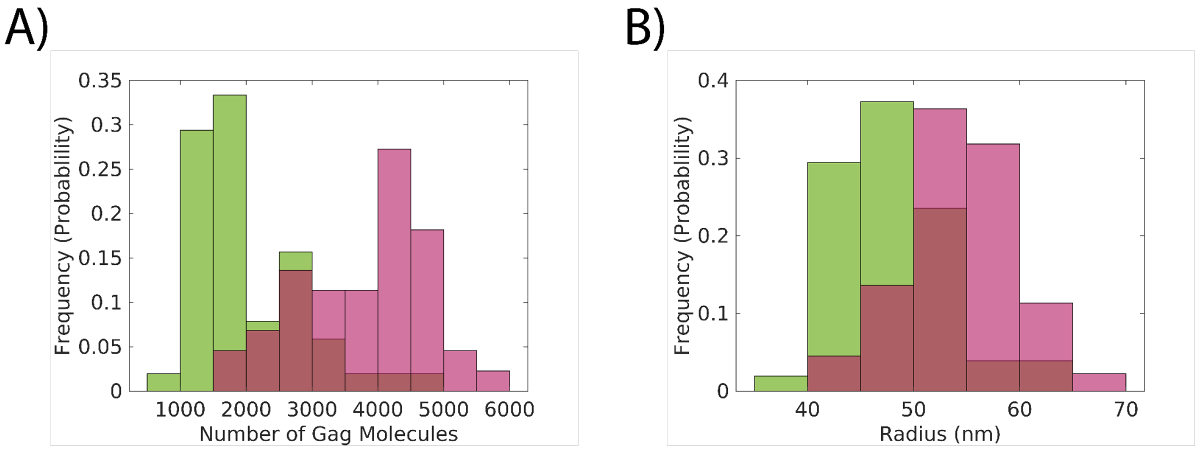


**Figure S4. (A)** Normalized histograms showing probability frequency for the number of Gag molecules in NL4.3:PR(D25N) immature virions (green) and humanized Gag VLPs (magenta). **(B)** Normalized histograms showing probability frequency for the membrane radius of NL4.3:PR(D25N) immature virions (green) and humanized Gag VLPs (magenta).

| Backbone Protein/Gene | Nucleotide Sequence |
|---|---|
| PR:(D25N) Mutagenesis Site | …GACCTACACCTGTCAACATAATTGGAAGAAATCTGTTGACTCAGATTGGCTGCACTTTAAATTTTCCC<br>ATTAGTCCTATTGAGACTGTACCAGTAAAATTAAAGCCAGGAATGGATGGCCCAAAAGTTAAACAATGG<br>CCATTGACAGAAGAAAAAATAAAAGCATTAGTAGAAATTTGTACAGAAATGGAAAAGGAAGGAAAAAT<br>TTCAAAAATTGGGCCTGAAAATCCATACAATACTCCAGTATTTGCCATAAAGAAAAAAGACAGTACTAA<br>ATGGAGAAAATTAGTAGATTTCAGAGAACTTAATAAGAGAACTCAAGATTTCTGGGAAGTTCAATTAGG<br>AATACCACATCCTGCAGGGTTAAAACAGAAAAAATCAGTAACAGTACTGGATGTGGGCGATGCATATTT<br>TTCAGTTCCCTTAGATAAAGACTTCAGGAAGTATACTGCATTTACCATACCTAGTATAAACAATGAGAC<br>ACCAGGGATTAGATATCAGTACAATGTGCTTCCAC… |
| Humanized Gag Sequence | CATGGGCGCCCGCGCCTCCGTGCTGTCCGGCGGCGAGCTGGACAAGTGGGAGAAGATCCGCCTGCGCCC<br>CGGCGGCAAGAAGCAGTACAAGCTGAAGCACATCGTGTGGGCCTCCCGCGAGCTGGAGCGCTTCGCCG<br>TGAACCCCGGCCTGCTGGAGACCTCCGAGGGCTGCCGCCAGATCCTGGGCCAGCTGCAGCCCTCCCTGC<br>AAACCGGCTCCGAGGAGCTGCGCTCCCTGTACAACACCATCGCCGTCCTGTACTGCGTGCACCAGCGCA<br>TCGACGTGAAGGACACCAAGGAGGCCCTGGACAAGATCGAGGAGGAGCAGAACAAGTCCAAGAAGAA<br>GGCCCAGCAGGCCGCCGCCGACACCGGCAACAACTCCCAGGTGTCCCAGAACTACCCCATCGTGCAGA<br>ACCTGCAGGGCCAGATGGTGCACCAGGCCATCTCCCCCCGCACCCTGAACGCCTGGGTGAAGGTGGTGG<br>AGGAGAAGGCCTTCTCCCCCGAAGTCATCCCCATGTTCTCCGCCCTGTCCGAGGGCGCCACCCCCCAGG<br>ACCTGAACACCATGCTGAACACCGTGGGCGGCCACCAGGCCGCCATGCAGATGCTGAAGGAGACCATC<br>AACGAGGAGGCCGCCGAGTGGGACCGCCTGCACCCCGTGCACGCCGGCCCCATCGCCCCCGGCCAGAT<br>GCGCGAGCCCCGCGGCTCCGACATCGCCGGCACCACCTCCACCCTGCAAGAGCAGATCGGCTGGATGAC<br>CCACAACCCCCCCATCCCCGTGGGCGAGATCTACAAGCGCTGGATCATCCTGGGCCTGAACAAGATCGT<br>GCGCATGTACTCCCCCACCTCCATCCTGGACATCCGCCAGGGCCCCAAGGAGCCCTTCCGCGACTACGT<br>GGACCGCTTCTACAAGACCCTGCGCGCCGAGCAGGCCTCCCAGGAGGTAAAGAACTGGATGACCGAGA<br>CCCTGCTGGTGCAGAACGCCAACCCCGACTGCAAGACCATCCTGAAGGCCCTGGGCCCCGGCGCCACCC<br>TGGAGGAGATGATGACCGCCTGCCAGGGCGTGGGCGGCCCCGGCCACAAGGCCCGCGTGCTGGCCGAG<br>GCCATGTCCCAAGTCACCAACCCCGCCACCATCATGATCCAGAAGGGCAACTTCCGCAACCAGCGCAAG<br>ACCGTGAAGTGCTTCAACTGCGGCAAGGAGGGCCACATCGCCAAGAACTGCCGCGCCCCCCGCAAGAA<br>GGGCTGCTGGAAGTGCGGCAAGGAGGGCCACCAGATGAAAGATTGTACTGAGAGACAGGCTAATTTTT<br>TAGGGAAGATCTGGCCTTCCCACAAGGGAAGGCCAGGGAATTTTCTTCAGAGCAGACCAGAGCCAACA<br>GCCCCACCAGAAGAGAGCTTCAGGTTTGGGGAAGAGACAACAACTCCCTCTCAGAAGCAGGAGCCGAT |

| | |
|---|---|
| | AGACAAGGAACTGTATCCTTTAGCTTCCCTCAGATCACTCTTTGGCAGCGACCCCTCGTCACAATAA |
| Ψ:(Δ 105-278 & Δ 301-332) Deletion Sites | TGTGCCCGTCTGTTGTGTGACTCTGGTAACTAGAGATCCCTCAGACCCTTTTAGTCAGTGTGGAAAATCT<br>CTAGCAGTGGCGCCCGAACAGGGACTTGAAAGCGAAAGTAAAGCCAGAGGAGATCTCTCGACGCAGGA<br>CTCGGCTTGCTGAAGCGCGCACGGCAAGAGGCGAGGGGCGGCGACTGGTGAGTACGCCAAAAATTTTG<br>ACTAGCGGAGGCTAGAAGGAGAGAG |
| Gag-Pol SS2-Mutagenesis Sites | …TAAAAGCATTGGGACCAGGAGCGACACTAGAAGAAATGATGACAGCATGTCAGGGAGTGGGGGGAC<br>CCGGCCATAAAGCAAGAGTTTTGGCTGAAGCAATGAGCCAAGTAACAAATCCAGCTACCATAATGATAC<br>AGAAAGGCAATTTTAGGAACCAAAGAAAGACTGTTAAGTGTTTCAATTGTGGCAAAGAAGGGCACATA<br>GCCAAAAATTGCAGGGCCCCTAGGAAAAAGGGCTGTTGGAAATGTGGAAAGGAAGGACACCAAATGAA<br>AGATTGTACTGAGAGACAGGCTAACTTCTTAGGGAAGATCTGGCCTTCCCACAAGGGAAGGCCAGGGA<br>ACTTCCTTCAGAGCAGACCAGAGCCAACAGCCCCACCAGAAGAGAGCTTCAGGTTTGGGGAAGAGACA<br>ACAACTCCCTCTCAGAAGCAGGAGCCGATAGACAAGGAACTGTATCCTTTAGCTTCCCTCAGATCACTC<br>TTTGGCAGCGACCCCTCGTCACAATAAAGATAGGGGGGCAATTAAAGGAAGCTCTATTAGATACAGGA<br>GCAGATGATACAGTATTAGAAGAAATGAATTTGCCAGGAAGATGGAAACCAAAAATGATAGGGGGAAT<br>TGGAGGTTTTATCAAAGTA… |
| ΔENV Mutagenesis Site | …AGTGCTACAGAAAAATTGTGGGTCACAGTCTATTATGGGGTACCTGTGTGGAAGGAAGCAACCACCA<br>CTCTATTTTGTGCATCAGATGCTAAAGCATATATGATACAGAGGTACATAATGTTTGGGCCACACATGCC<br>TGTGTACCCACAGACCCCAACCCACAAGAAGTAGTATTGGT… |

**Table S5:** Nucleotide sequences for mutagenesis sites of NL4.3 and humanized Gag plasmids presented. Green highlighting identifies mutated nucleotides, red identifies deleted nucleotides, and blue identifies added nucleotides.

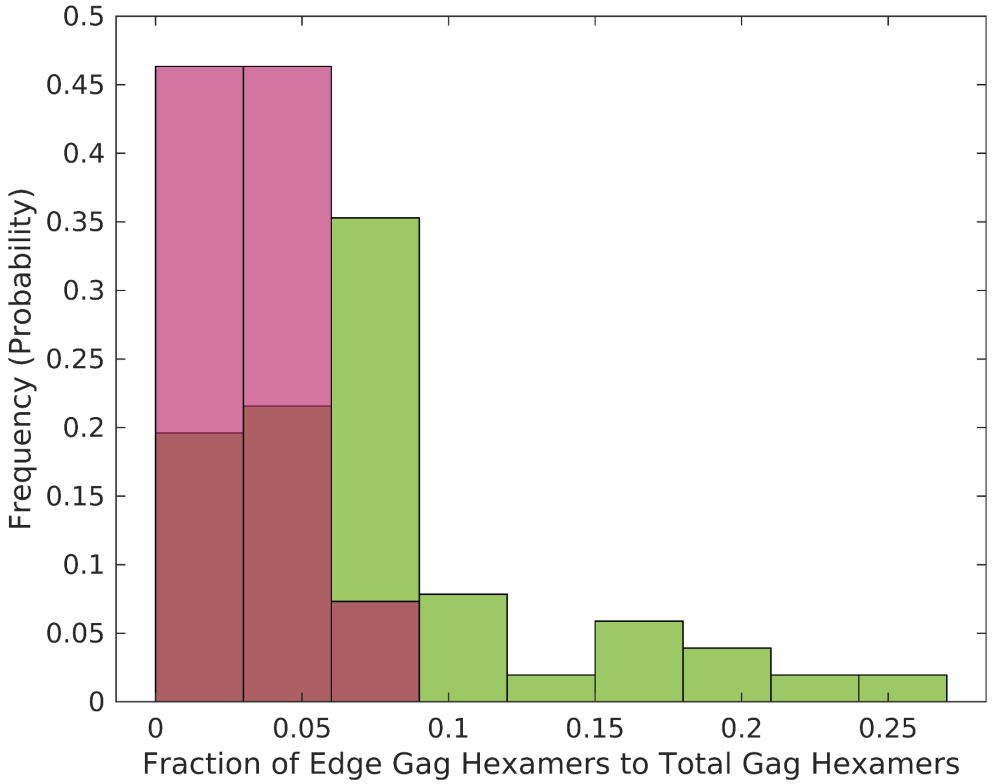

**Figure S6:** Normalized histogram showing ratios of edge Gag hexamers to total Gag Hexamers compared between NL4.3:PR(D25N) immature virions (green) and humanized Gag VLPs (magenta).
